# Effects of Exogenous Nitric Oxide Gas on *Mycobacterium tuberculosis* in vitro and in mice

**DOI:** 10.64898/2026.08.18.744881

**Authors:** Xiuju Jiang, Carl Nathan

## Abstract

In 1992, inhaled NO (iNO) at low doses entered the practice of medicine for cardiopulmonary indications. Recently, iNO at higher doses has been tested in diverse pulmonary infections. However, nothing is known about the ability of exogenous NO gas to kill *Mycobacterium tuberculosis* (Mtb), the leading cause of death from infection between major viral pandemics. Here we mimicked exposure conditions used in recent human studies of high-dose iNO to explore the effects of NO gas against Mtb in vitro and in mice. We saw a profound bactericidal effect of NO gas in vitro against Mtb incubated in shallow, mildly acidic fluid. Mtb-infected mice tolerated inhaled NO well, except for developing more methemoglobinemia than humans at the same level of exposure. In Mtb-infected mice with poorly aerated pulmonary infiltrates, inhaled NO had an anti-inflammatory effect but did not reduce the bacterial burden. These results may help inform the decision whether to test inhaled NO as an adjunctive treatment for tuberculosis, and if so, in what settings and with what goals.

## INTRODUCTION

As recently reviewed (Nathan, 2026), endogenously produced NO has a physiological role as an anti-infective agent. This has encouraged investigators to explore the delivery of exogenous NO to sites of infection, using topical and systemically administered agents that generate NO itself or NO-related moieties, that is, products of NO’s reaction with oxygen that remain chemically reactive (reactive nitrogen intermediates; RNS) and products of the reduction of nitro groups on organic compounds that can bond with biological molecules.

Another way to deliver exogenous NO is by inhalation. Inhaled NO (iNO) entered the practice of medicine in 1992 with the treatment of infants suffering from persistent pulmonary hypertension (Kinsella et al., 1992; Roberts et al., 1992). In 1999, the US Food and Drug Administration approved NO gas as a drug (Food and Drug Administration, 1999). Nonetheless, as a potential anti-infective agent, iNO faces challenges (Nathan, 2026). Among these is NO’s rapid reaction with oxygen to form a more toxic radical, ^•^NO_2_. Another challenge has been the provision of NO by chemical supply companies in tanks containing NO at levels far above what people can safely inhale. Until recently, such challenges restricted the administration of iNO to medical facilities with equipment that can deliver NO at levels that keep methemoglobin at < ∼ 5% while keeping ^•^NO_2_ < ∼ 5 ppm. Recently, portable equipment has become available that uses an electric pulse and a catalyst to generate NO continuously from oxygen and nitrogen in ambient air at the point of care, scrub resultant NO_2_ and deliver NO doses up to 300 ppm for inhalation. That engineering breakthrough encouraged us to use such equipment and experimental dosing regimens from studies with large animals and humans (Yu et al., 2026) to study the impact of exogenous NO gas on Mtb in vitro and in mice.

## RESULTS

### Antimycobacterial effects of intermittent exposure to NO gas in vitro

The ability of RNS to inhibit or kill Mtb in vitro has only been demonstrated with continuous exposure, whether to RNS-generating acidified NO_2_^-^, NO-donating NONOates, or iNOS-expressing macrophages (Ehrt et al., 2001). In contrast, when patients inhale NO gas at high doses, the inhalations typically last 30 min and may be repeated at widely spaced intervals.

Before treating mice in a similar manner, we asked three questions. Is intermittent exposure to NO gas able to kill Mtb in vitro under conditions that avoid build-up of ^•^NO_2_? Does intermittent exposure to NO avoid the induction of phenotypic tolerance to TB drugs that is seen with continuous exposure to RNS (Liu et al., 2016; Voskuil et al., 2003)? Can intermittent exposure to NO gas synergize with compounds whose antimycobacterial action is RNS-dependent under conditions of continuous exposure to RNS (Warrier et al., 2015; Warrier et al., 2026)?

We decided to conduct in vitro experiments under the same conditions as we would subsequently use to treat mice. Rather than force immobilized mice to breathe from nosecones, we allowed them free movement in a Lucite chamber. We used the same chamber for experiments in microtest plates. We began these studies with a custom-built NO generator from Third Pole Therapeutics (Waltham, MA) (**Supplemental Figure 1A, B)** and switched to a prototype of a commercial unit after Third Pole Therapeutics modified it to produce the same dose of NO (300 ppm) used by Yu et al. (Yu et al., 2026) (**Supplemental Figure 1C)**. We flowed air through the chamber at a rate that kept ^•^NO_2_ < 5 ppm.

We began by testing the effect of 30-min exposures of virulent Mtb H37Rv to 300 ppm NO under diverse conditions: in 24-, 96- and 384-well plates with volumes of medium ranging from 0.5 mL to 0.05 mL to vary the diffusion distance from the gas phase through the liquid phase. We compared results in PBS and 7H9 medium at pH 6.8. We also tested 7H9 at pH 5.5 with the thought that acidic pH promotes recycling of nitrite (a product of NO’s oxidation in air) to NO. Indeed, acid-catalyzed recycling of nitrite to NO may contribute to the anti-mycobacterial of IFN-γ-activated macrophages, which acidify their phagolysosomes (Schaible et al., 1998). We compared the impact of 4-6 cycles of NO per day over 2-4 days, resulting in exposures ranging from 1200 to 3600 ppm⋅h. As a control, Mtb was exposed under the same conditions in the same device to the same flow of air without NO.

Cyclic exposure to NO over 4 days totaling 3600 ppm⋅h led to a ∼10-fold reduction of Mtb CFU when the Mtb was incubated in 7H9 medium at pH 5.5 in 100 μL volumes in 96-well plates (fluid height, 2.49 mm) (**Figure 1A**). By comparison to the other conditions tested, where less killing was seen, it appeared that the key features permitting bactericidal activity of NO gas above a liquid culture of Mtb were shallow fluid depth and mildly acidic pH. The dramatic impact of fluid depth was confirmed in direct tests (**Figure 1B**). In wells with 60 μL of fluid, the reduction in CFU was >2 log_10_, while at 50 μL and below it was ∼6 log_10_ to below the limit of detection (2 CFU).

**Figure 1.**
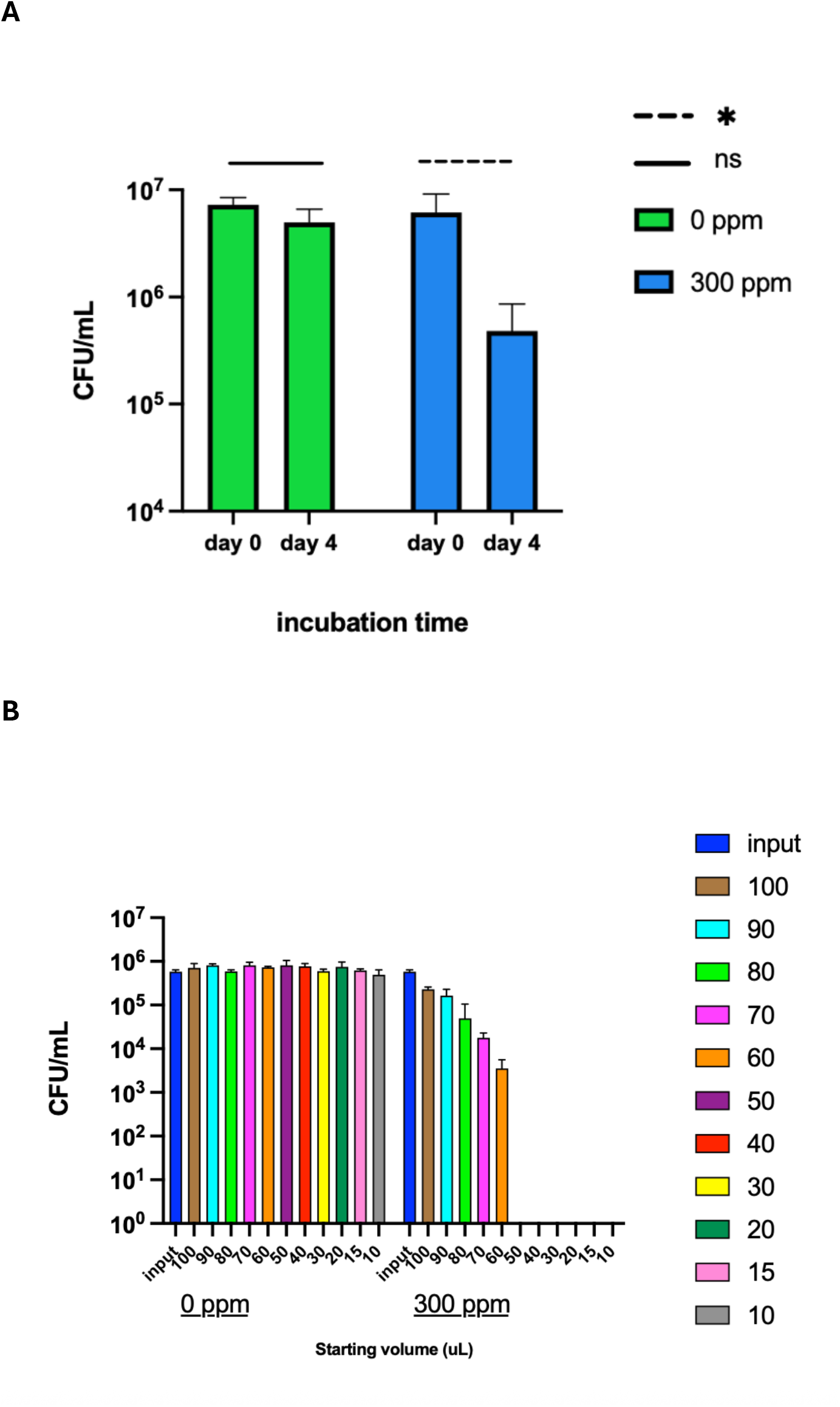
Bactericidal action of NO gas against Mtb in vitro. **(A)** Results in wells with a fluid height of 2.49 mm. Data are means ± SD from 4 independent experiments, each in triplicate, where the mean shown is the mean of the means from each experiment and the SD shown is the mean of the SDs from each of the experiments. Mtb H37Rv was incubated in 96-well plates in 100 μL 7H9 medium adjusted to pH 5.5 with MES. Plates were exposed to air containing 0 or 300 ppm NO for 30 min per exposure for 6 exposures per day over 4 days (cumulative exposure, 3600 ppm⋅h). Day 0 values are CFU/mL before any exposure. Green bars, NO = 0 ppm. Blue bars, NO = 300 ppm. **(B)** As in **(A)** but the same number of Mtb was plated in wells with the volumes of fluid indicated in μL. Results are means ± SD of triplicates in one of 2 experiments with similar results. Solid horizontal lines indicate non-significant differences; dashed lines, p < 0.05 by Student’s t test.

Cyclic exposure to NO did not impair the in vitro mycobactericidal action of rifampin. This was the case whether the depth of fluid in a 24-well plate precluded a direct mycobactericidal action of NO gas (**Figure 2A**) or its shallowness in a 96-well plate permitted NO gas to kill some of the Mtb on its own (**Figure 2B**).

**Figure 2.**
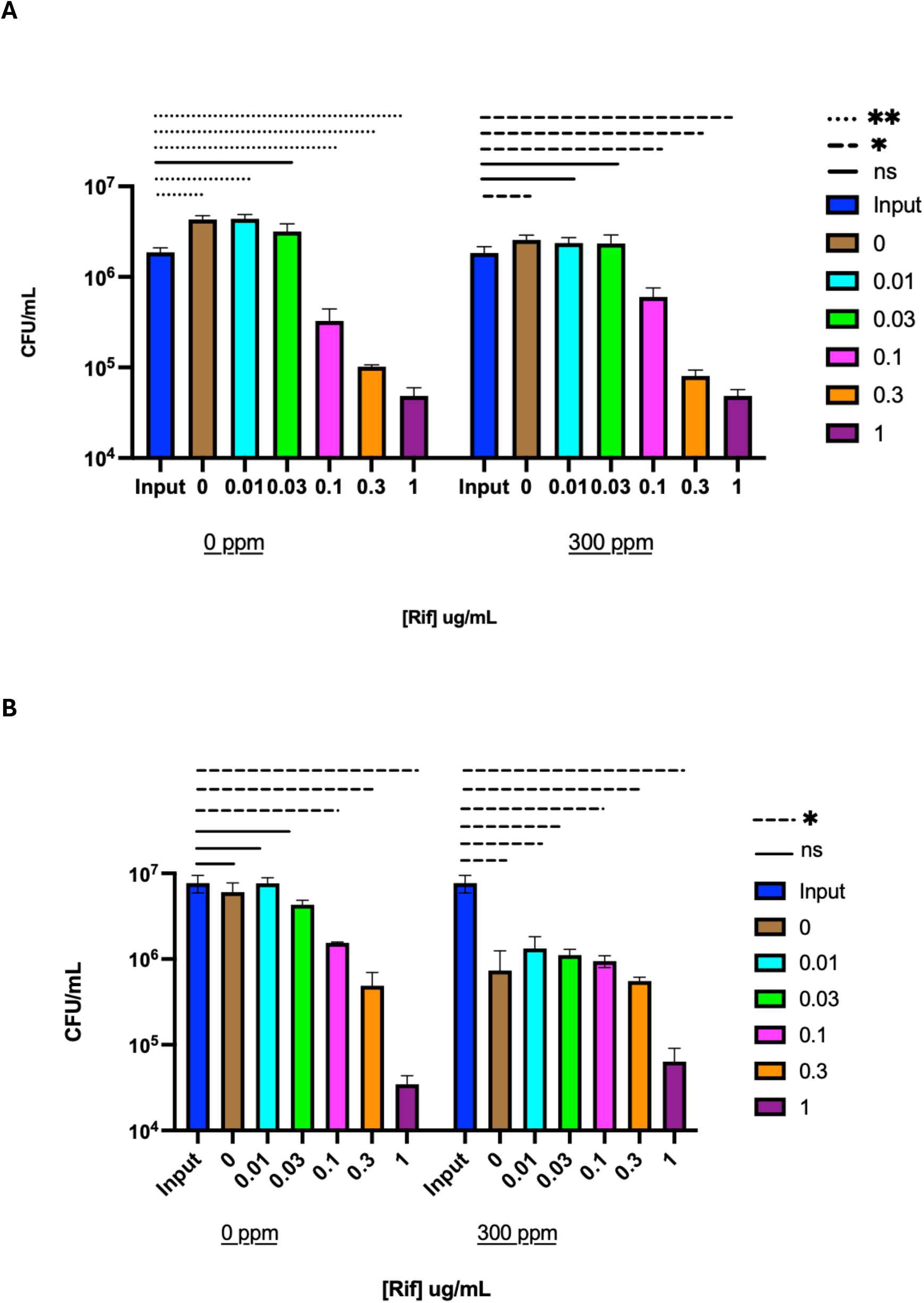
Non-interference in mycobactericidal action of rifampin by cyclic exposure of Mtb to NO gas in vitro. **(A)** Under conditions where NO gas alone was ineffective (in 500 μL 7H9 medium, pH 6.8, in a 24-well plate, giving a fluid height of 420 mm). **(B)** Under conditions where NO gas alone was effective (in 100 μL 7H9 medium, pH 5.5, in a 96-well plate, giving a fluid height of 2.49 mm.) Data in both **(A)** and **(B)** are means ± SD from one experiment representative of 2 independent experiments of each type, each in triplicate. Solid horizontal lines indicate non-significant differences; dashed lines, p < 0.05; dotted lines, p < 0.01 by Student’s t test.

On the contrary, cyclic exposure to NO synergized with two chemically distinct compounds— a bromoindazole (Warrier et al., 2015) (**Figure 3A**) and a diarylindole (Warrier et al., 2026) (**Figure 3B)**— whose rapid and extensive antimycobacterial action is dependent on RNS under conditions where a sustained flux of RNS is generated by dismutation of acidified nitrite or by decomposition of an NO-donating compound. The synergy was seen under conditions where NO alone was inactive and where the bromoindazole and the diarylindole alone were inactive. Together, the compounds and cyclic exposure of Mtb to NO gas produced a ∼4 log_10_ drop in CFU within 4 days.

**Figure 3.**
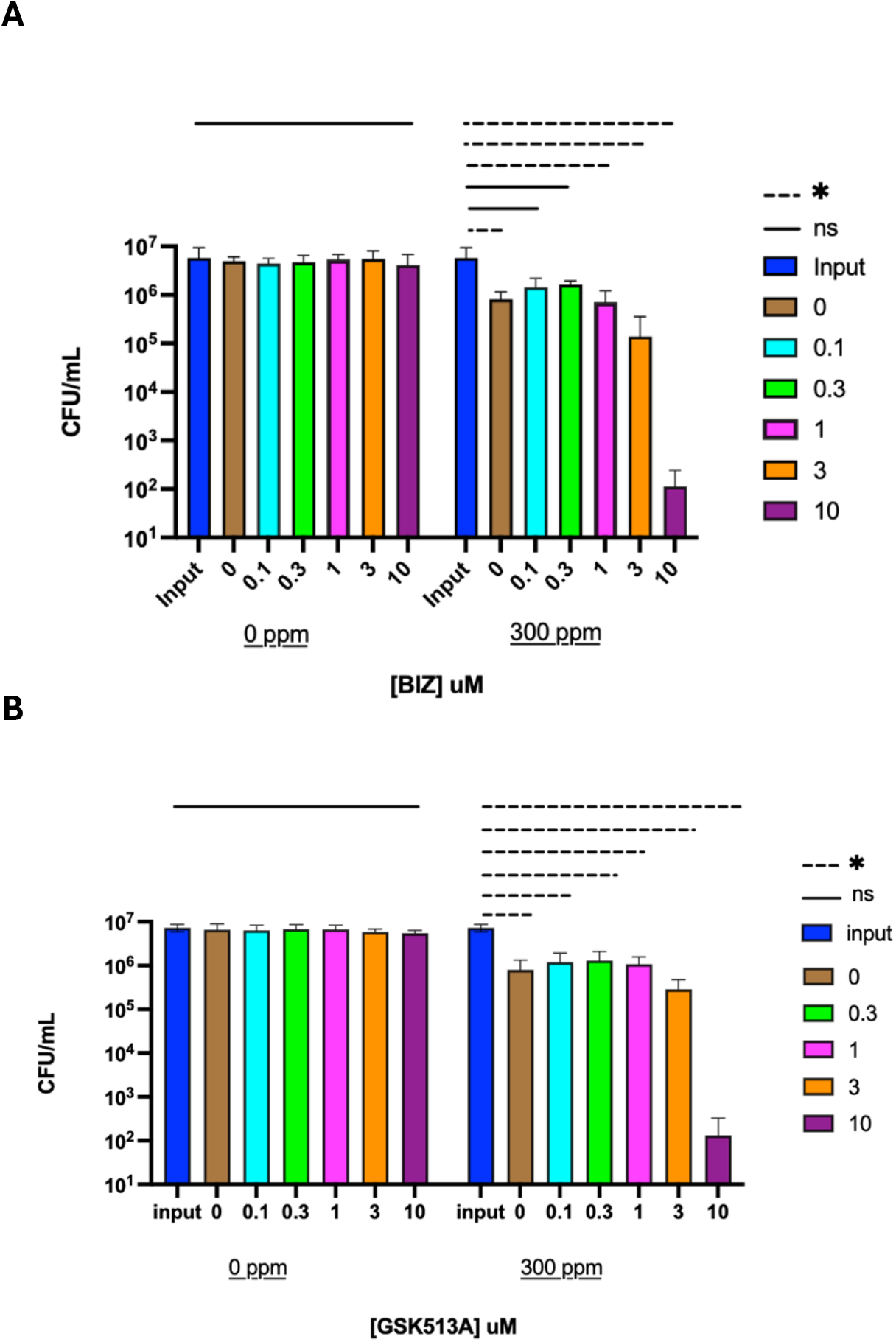
Augmentation of mycobactericidal action of (A) bromoindazole 12 (Warrier et al., 2015) and (B) diaryl-aminoindazole GSK1867513A (GSK513A) (Warrier et al., 2026) by cyclic exposure of Mtb to NO gas in vitro. In both **(A)** and **(B)** the data are means ± SD from 2 independent experiments, each in triplicate, where the mean shown is the mean of the means from each experiment and the SD shown is the mean of the SDs from each of the experiments. Exposure to NO was as described in Figure 1 (cumulative exposure, 3600 ppm⋅h). Solid horizontal lines indicate non-significant differences; dashed lines, p < 0.05 by Student’s t test.

We tried to determine if NO gas would affect Mtb within macrophages, using PMA-differentiated human THP1 cells. Unfortunately, the high rate of air flow in the exposure chamber was toxic to the monolayers, both with and without NO in the airstream, despite replacement of evaporative losses at intervals. Thus, these results were uninterpretible.

### Tolerability of iNO in mice

To test iNO in mice, we chose a dose of 300 ppm NO, because 150-350 ppm, often given over 30 min, were the highest doses reported for use in humans at the time we initiated these studies (Bartley et al., 2020; Safaee Fakhr et al., 2021; Strickland et al., 2022; Wiegand et al., 2020). As a control, mice housed in the same chamber received NO-free air at the same flow rate. When returned to their cages after exposure, Mtb-infected mice immediately moved about rapidly. In one early experiment, two mice that had not been infected with Mtb but were recorded as being in poor health died promptly after NO exposure. Thereafter, over the course of 9 experiments with 65 Mtb-infected mice inhaling NO up to 15,750 ppm⋅h over periods ranging from 4 to 15 days, there were no signs of morbidity and no deaths.

Clinical administration of iNO is monitored by the formation of methemoglobin, which is generally held below 5-6% and quickly falls near or below 1% when inhalation of NO is discontinued. A method for measuring methemoglobin in a drop of blood that was recommended for small animals (Valsecchi et al., 2022) gave erratic results in our hands (**Supplemental Fig. 2A, 2B)**. Accordingly, we resorted to a monitor used in pediatric hospital units that measures oxygen saturation, methemoglobin, pH, electrolytes and metabolites in small blood volumes, with the modifications described in Methods to deal with the even smaller volumes available from mouse tail vein nicks. Pre-exposure methemoglobin levels were 0.3% ± 0.3 (mean ± SD) in 3 experiments. Post-exposure levels averaged 49.5 ± 1.9 % (mean ± SD) in 6 experiments (**Figure 4**), which we subsequently realized confirmed the observation of Wiegand et al. with mice breathing 300 ppm of NO for 30 min (Wiegand et al., 2021). When some of the mice were tested again a day later, they developed the same level of methemoglobin (**Figure 4**), indicating an inability to adaptively increase methemoglobin reductase activity. Such levels of methemoglobin in humans cause dyspnea, headache and fatigue (Rockwood et al., 2003). Methemoglobin levels returned to baseline in mice breathing room air with a half-time of 30 min (**Figure 4**). In contrast, peak methemoglobin averaged 5.5% in 10 non-exercising human volunteers breathing NO at 300 ppm for 30 min in 148 sessions (Yu et al., 2026).

**Figure 4.**
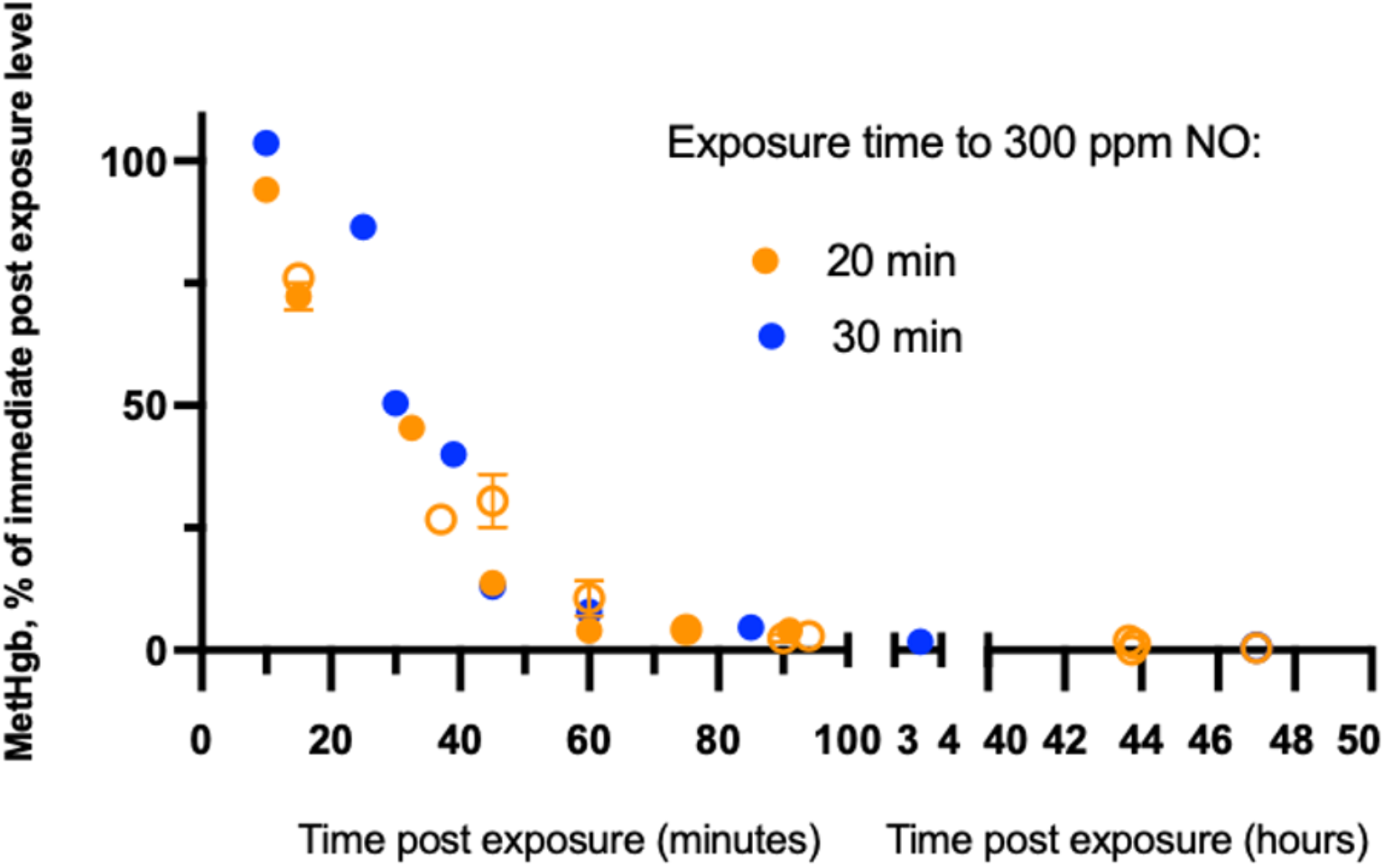
Methemoglobin levels measured by clinical analyzer in mice after a 20- or 30-minute exposure to NO at 300 ppm and rate of decline in methemoglobin levels upon inhalation of room air (0 ppm NO). Symbols without error bars represent results from individual mice. Symbols with error bars represent means ± SDs for results from 3-5 mice at the indicated time points. Closed symbols represent results from mice tested once. Open symbols represent results from some of those mice that were tested again the following day, indicating that there had been no induction of resistance to methemoglobinemia. Pre-exposure methemoglobin levels were 0.3% ± 0.3 (mean ± SD) in 3 experiments. Post-exposure levels averaged 49.5 ± 1.9 % (mean ± SD) in 6 experiments.

The peak level of methemoglobin we observed in mice would be unacceptable in clinical practice. However, given that no such elevations are seen with the equivalent exposure in humans and that Mtb-infected mice showed no adverse effects on motility, survival or blood pH, [Na^+^], [K^+^], [Ca^++^], [Cl^-^], [glucose] or [lactate] (not shown), we proceeded to treat Mtb-infected mice with 30-min exposures, with rests in room air for ≥ 60 min between exposures.

### Effects of iNO in mice with tuberculous pneumonia

Giving Mtb-infected mice multiple exposures to NO per day over multiple days afforded the opportunity to monitor the following additional indicators of tolerability: body weight, lung weight, liver function tests and histopathology. As shown in **Table 1**, lack of adverse effects on body weight and liver function further supported that iNO was well tolerated, aside from methemoglobinemia. Lack of a differential increase in the weight of the lungs suggested that there was no greater accumulation of edema or of infiltrating cells than in the control group.

**Table 1.** Evidence for the nontoxicity of NO inhalation by Mtb-infected mice. Cumulative exposures are indicated for the groups treated with NO at 300 ppm x 30 min for 4-7 times per day for 4-10 days over 1-2 weeks, omitting weekends. The cumulative NO exposure is shown for the group that received NO. In each experiment, a control group was handled exactly the same except that NO in the exposure chamber in-flow was at 0 ppm. Treatment was started at the indicated number of days after inhaling Mtb H37Rv sufficient to give ∼100 CFU per lung as measured the following day (called day 0). There were 5 mice in each group. Numbers for weights are means ± SD.

| Experiment | 11 | 14 | 13 | 16 | 22 | 24 | 30 |
| --- | --- | --- | --- | --- | --- | --- | --- |
| Day start<br>iNO<br>(duration) | 7 (4 d) | 7 (5 d) | 28 (4 d) | 28 (5 d) | 28 (9 d) | 28 (5 d) | 42 (10 d) |
| Exposure<br>(ppm·hr) | 1200 | 3900 | 1950 | 4950 | 9450 | 5250 | 10,500 |
| Deaths | 0 | 0 | 0 | 0 | 0 | 0 | 0 |
| Motility | WNL | WNL | WNL | WNL | WNL | WNL | WNL |
| Body<br>weight, day<br>of starting<br>Rx (g) | 20.36 ±<br>0.93 | 18.45 ±<br>0.84 | 20.42 ±<br>1.67 | 20.37 ±<br>1.98 | 19.73 ±<br>1.62 | 20.41 ±<br>0.99 | 20.88 ±<br>1.13 |
| Body<br>weight,<br>change<br>from day 1<br>for NO<br>group (g) | +0.68 | +0.63 | +0.90 | +2.04 | +2.42 | +1.77 | +2.40 |
| Body<br>weight,<br>change<br>from day 1<br>for control<br>group (g) | +0.72 | +0.46 | +0.91 | +2.04 | +1.99 | +1.23 | +1.30 |
| Lung<br>weight day<br>of starting<br>Rx (g) | 0.22 ±<br>0.03 | 0.21 ±<br>0.04 | 0.31 ±<br>0.06 | 0.27 ±<br>0.03 | 0.27 ±<br>0.02 | 0.36 ±<br>0.04 | 0.22 ±<br>0.05 |
| Lung weight at sac for NO group (g) | 0.26 ± 0.04 | 0.22 ± 0.03 | 0.33 ± 0.03 | 0.31 ± 0.02 | 0.31 ± 0.03 | 0.38 ± 0.05 | 0.27 ± 0.04 |
| Lung weight at sac for control group (g) | 0.23 ± 0.02 | 0.21 ± 0.03 | 0.33 ± 0.04 | 0.34 ± 0.06 | 0.31 ± 0.03 | 0.40 ± 0.06 | 0.28 ± 0.04 |
| LFTs plus | WNL | WNL | WNL | WNL | WNL | pending | WNL except note 1 |

Abbreviations: iNO, inhaled NO. LFTs plus: liver function tests (alkaline phosphatase, alanine transaminase, aspartate transaminase, $\gamma$ -glutamyl transferase, bilirubin) plus total serum protein, albumin, globulin, cholesterol, triglycerides. WNL, within normal limits.
Note 1: All groups in this study had normal values for all tests except aspartate transaminase levels, which were above the normal upper limit (77 U/L) from day 1 onward with no change.

Likewise, scoring by pathologists who were blinded to the treatment groups indicated that exposures to iNO of 4950 and 9450 ppm⋅hr over 5 to 9 days reduced the composite pathology score and the degree of inflammation, without exacerbating consolidation, edema formation, fibrin deposition, or hemorrhage, compared to mice undergoing the same treatments without NO in the airstream (**Figure 5**).

**Figure 5.**
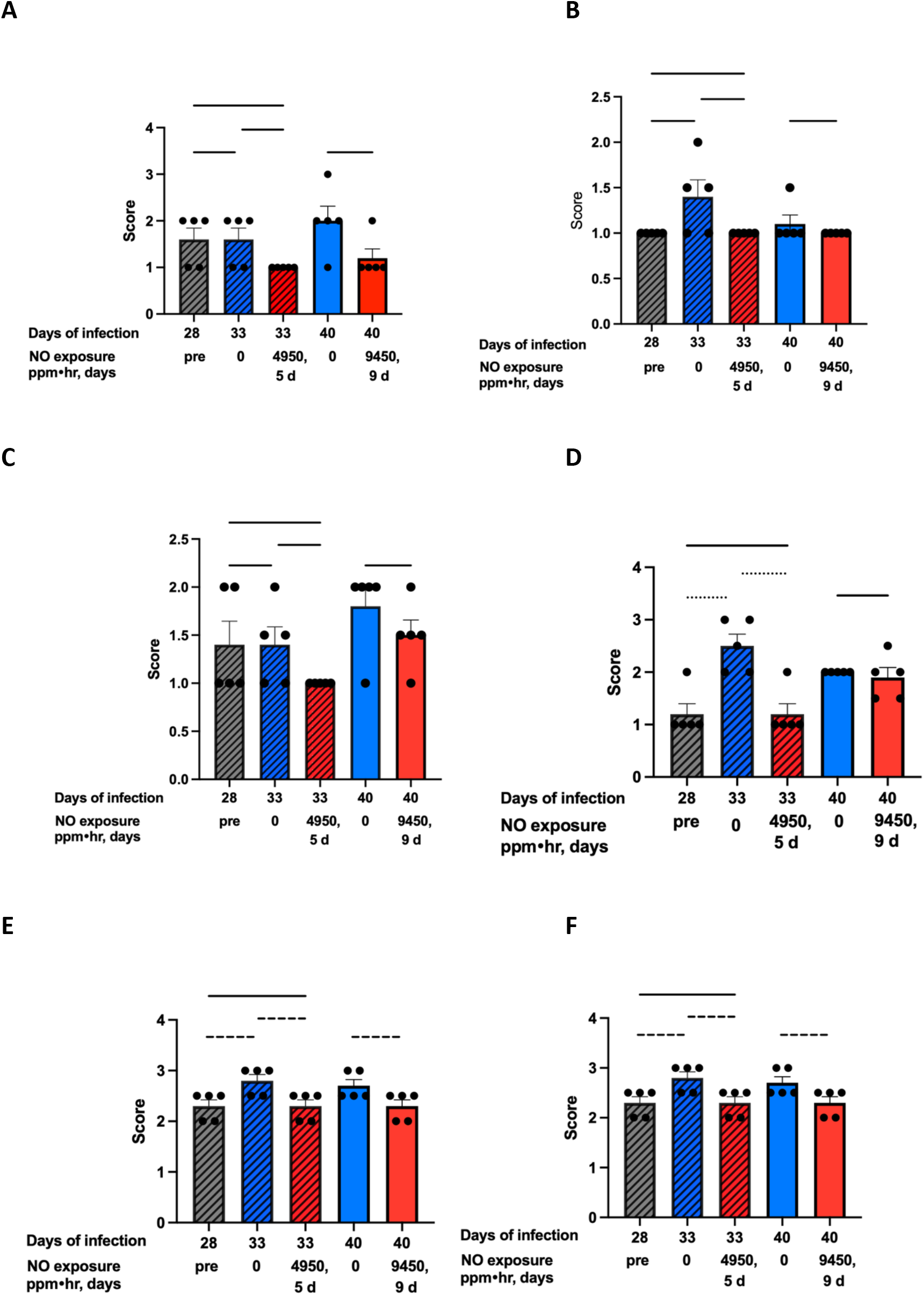
Non-inflammatory impact of inhaled NO on the histopathology in lungs of Mtb-infected mice. **(A)** Consolidation. **(B)** Edema. **(C)** Fibrin deposition. **(D)** Hemorrhage. **(E)** Inflammation. **(F)** Composite pathology score. Results were recorded by two veterinary pathologists who were blind to the treatments. The first pathologist scored one hematoxylin- and eosin-stained slide from lungs of each of 5 mice in each of the two experiments shown, which involved cumulative exposures of 4950 and 9450 ppm⋅hr. The second pathologist found similar results in a 3^rd^ experiment in which the total exposure was 5250 ppm⋅hr. Scoring: 0, the feature was absent; 1, present in <25% of the field; 2, 26-50%; 3, 51-75%; 4, > 75%. Solid horizontal lines indicate non-significant differences; dashed line, p < 0.05; dotted line, p < 0.01 by Student’s t test.

In four experiments, we treated C57BL/6 mice beginning on day 28 or later after aerosol infection with an Mtb H37Rv strain positive for the virulence factor phthiocerol dimycocerosate (PDIM) so as to deposit ∼100 CFU per lung, as measured the following day (day 1). By day 28, pneumonitic infiltrates were well established. We saw a 63% reduction in CFU in one experiment with an exposure of 9450 ppm⋅h but no reduction in the three other experiments with exposures of 5250, 10,500 and 15,750 ppm⋅h, respectively. When mice were co-treated with iNO and rifampin, iNO did not impair the action of rifampin.

In sum, aside from methemoglobinemia, iNO under the conditions used appeared to be safe but ineffective in mice with tuberculous pneumonia.

## Discussion

In 1999, Long et al. (Long et al., 1999) reported exposing Mtb on agar plates in a large chamber to 90 ppm NO for 48 h, resulting in an exposure of 4320 ppm⋅h, with the result that CFU were reduced from ∼100 to 0. However, ^•^NO_2_ almost certainly formed under these conditions. To our knowledge, in the ensuing 27 years, there has been no further report of the effect of exogenous NO gas on Mtb. Thus, the ability of the less toxic NO itself to kill Mtb when delivered as a gas had apparently never been tested. We can now affirm that NO gas at 300 ppm flowing above a shallow bed of mildly acidified fluid for 30-minute periods repeated over several days can rapidly reduce the number of Mtb CFU in the fluid to below the limit of detection. Sublethal exposures to NO gas did not make Mtb tolerant of rifampin. However, NO gas did make Mtb highly susceptible to two compounds whose marked anti-mycobacterial activity was seen earlier to be strictly dependent on exposure to other NO-related moieties, which were delivered continuously rather than intermittently (Warrier et al., 2015; Warrier et al., 2026).

The exposure conditions that seemed optimal in vitro did not appear to pertain in C57BL/6 mice during the chronic phase of infection with Mtb H37Rv, because iNO had no consistent impact on the number of Mtb CFU recovered from their lungs. We speculate that pneumonitic pathology precluded iNO from reaching most of the bacteria, including the majority that reside within macrophages. While the C57BL/6 mouse model does not recapitulate key features of human TB pathology, such as formation of cavities, it did allow the demonstration that iNO as delivered here was well tolerated. The one exception was the marked susceptibility of the mice to accumulation of methemoglobin at levels not seen in humans inhaling NO at the same dose (Yu et al., 2026). Marked differences among species in their ability to reduce methemoglobin have been reported (Rockwood et al., 2003; Smith and Beutler, 1966).

The apparent utility of iNO in the treatment of non-tuberculous mycobacterial disease (Nathan, 1996) together with the mycobactericidal activity of NO gas in vitro and the tolerability of iNO in Mtb-infected mice described here may encourage consideration of iNO as an adjunctive treatment for people with TB. A reasonable aspiration would be to reduce viable Mtb in exhalations, given that exhalations come from sites at least partly accessible to inhalation and therefore to iNO. A secondary beneficial effect could be improvement of respiratory function by reduction of ventilation-perfusion mismatch.

## METHODS

### NO delivery

We used a Lucite exposure chamber measuring 16 cm x 10 cm x 5 cm (volume 760 cm^3^) to provide enough space either for microtest plates or for mice in groups of five while limiting excess airspace that could foster generation of ^•^NO_2_. When the chamber contained mice, we reduced airspace further by lining the chamber with a rubber pad 2.6 cm in height. We maintained the chamber in a biologic safety cabinet. Third Pole Therapeutics (Waltham, MA) generously shared a prototype of a high-dose, point-of-care NO generator for these studies. Because the mice were freely moving rather than breathing from nose cones, the device had to be adapted to minimize dwell time of NO in the chamber. **Supplemental Figure 1** schematizes the essential elements; **Supplemental Figure 2** provides photographs. The generator used electrical pulses acting on room air to produce a mix of NO and ^•^NO_2_ flowing at 8L/min. Passage through scrubbers (cylinders packed with calcium oxide, sodium hydroxide and potassium hydroxide) (Ishibe et al., 1995) removed the ^•^NO_2_ but left NO. Given that 2NO + O_2_ → 2^•^NO_2_ with a t_1/2_ = 27 min for 200 ppm NO in room air (Lundberg et al., 2008), we wanted to minimize ^•^NO_2_ formation by moving NO rapidly through the chamber to minimize its reaction time with O_2_. Accordingly, the output from the generator was merged with a HEPA-filtered air stream flowing at 17 liters per minute from a compressor. This succeeded in maintaining an atmosphere in the chamber in which NO was at 300 ppm while ^•^NO_2_ was 4.8 ppm. The combination of electrolysis in the generator and HEPA filtration of the compressor’s air flow sterilized the input gasses entering the chamber, preventing airborne contamination of the microtest plates, which we left open. Outflow from the chamber passed through a charcoal/potassium permanganate scrubber to remove both NO and ^•^NO_2_ and then through a HEPA filter to remove any Mtb before release into the biologic safety cabinet, whose own exhaust system provided further HEPA filtration. A sampling port allowing access to the chamber outflow passed through a HEPA filter to an analyzer that recorded the chamber’s NO and ^•^NO_2_ levels. The analyzer was calibrated by alternately testing against room air and against pure NO and pure ^•^NO_2_ from commercially sourced cylinders (Ideal Calibrations, Melvindale, MI 48122). Additional hand-held, battery-powered monitors for NO and ^•^NO_2_confirmed that neither gas was detectable in the vicinity of the operator.

Both for studies in vitro and in mice, experimental and control groups were alternated in the exposure chamber for every treatment cycle in every experiment.

For in vitro studies, a test tube rack in the exposure chamber held a microtest plate on each of two shelves. Given that 2NO + O_2_ → 2^•^NO_2_, and 2^•^NO_2_ + H_2_O → HNO_2_+ HNO_3_, we measured accumulating nitrite (NO_2_^-^) by the Griess reaction (Ding et al., 1988) as confirmation that each position in the microtest plate received an equivalent exposure to NO (data not shown). However, measurement of the volume in each well after a 30-min exposure indicated greater evaporation from the lower rack, which was directly opposite the inflow, than from the upper rack. Therefore, we used the lower microtest plate to hold water for humidification and used the upper plate for experiments. During experiments, we measured residual volume in sentinel wells in the upper plate after each 2-4 exposures and added sterile water in volumes equivalent to the evaporative loss.

### Mtb

Virulent, phthiocerol mycoserosate-expressing Mtb H37Rv was cultured under 5% CO_2_ in air in Middlebrook medium with 0.2% glycerol, 0.5% BSA, 2% dextrose, 0.85% NaCl, and 0.02% tyloxapol. CFU were counted 3 weeks after serial dilution in PBS with 0.02% tyloxapol and plating on 7H10 agar.

### NO exposure of Mtb in vitro

When indicated, 7H9 medium was buffered to pH 5.5 with 100 mM 2-(N-morpholino)ethanesulfonic acid hydrate (MES) (Sigma-Aldrich). Serial dilutions were made in PBS with 0.02% tyloxapol for plating on 7H10 agar to count CFU 3 weeks later.

### Infection of mice

Mice were studied under a protocol approved by the Institutional Animal Care and Use Committee of Weill Cornell Medicine. C57BL/6 female mice aged 8-10 weeks from Jackson Laboratories (Bar Harbor, ME) were rested 1-2 weeks, weighed and infected with Mtb by inhalation as described (Darwin and Nathan, 2005) to deliver ∼100 CFU per lung.

Mice were randomly assigned to control or treatment groups. Mean body weights in each group were indistinguishable before the onset of the experiments. At indicated times, mice were weighed, then euthanized by CO_2_ inhalation. Cardiac blood was collected and the serum was sterile-filtered for liver function tests in the clinical laboratory of the Research Animal Resource Center of Weill Cornell Medicine and Sloan Kettering Institute. The entire lung was weighed and then the left upper lobe was excised. All lung lobes except for the left upper lobe were processed for CFU, while the left upper lobe was fixed in formalin for histology as described (Darwin and Nathan, 2005).

### Methemoglobin measurements

To use the blood spot colorimetric method recommended for studies on small animals (Patton et al., 2016; Shihana et al., 2010), we constructed a standard curve by determining the minimal amount of NaNO_2_ that would convert all the hemoglobin in a sample of human blood to methemoglobin (Smith and Beutler, 1966).

For this, we used a 1:1 dilution of blood with distilled water and made it 10 mM in NaNO_2_ by adding 5% by volume of 200 mM NaNO_2_. This was designated as 100% methemoglobin and used for dilutions with various proportions of the 1:1 dilution of blood with distilled water.

Diluting each sample 1:3 with water improved sensitivity. We spotted 7.5 μL of each dilution on Whatman #1 filter paper, scanned the paper in the red channel on an Epson scanner and used ImageJ to score intensity per pixel in the central portion of the spots. We constructed standard curves with mouse blood in the same manner.

After determining that the filter spot method was imprecise, we rented a Radiometer ABL90 clinical blood monitor and operated it as instructed by the manufacturer (Radiometer Medical, Brea, CA). The device requires 65 μL of blood per sample, which is too much to get from mouse tail snips. We determined that diluting 3 ∼6-μL drops of tail vein blood in 70 μL of heparinized PBS did not reduce the level of hemoglobin below the Radiometer’s range of reliable quantification (≥ ∼2.5 g/dL) and allowed the instrument to report the percentage of methemoglobin as well as pH, [Na^+^], [K^+^], [Ca^++^], [Cl^-^], [glucose] and [lactate]. Gas sterilization would destroy the sensor mechanism in the Radiometer, so we were not able to bring it into the BSL3 facility and thus were only able to measure methemoglobinemia in mice that were not Mtb-infected.

### Other reagents

Bromoindazole (compound **12** in (Warrier et al., 2015)) was a kind gift of Alfonso Mendoza-Losano, GSK, Tres Cantos, Spain. The diarylindazole GSK1867513A, originally synthesized at GSK, Tres Cantos Spain, was resynthesized for this study by Kelin Li and Jeffrey Aubé (University of North Carolina), as reported (Warrier et al., 2026).

## ACKNOWLEDGMENTS

We thank Basil Athenson, Greg Hall, Jonathan Schiff and Ziad Elghazzawi (Third Pole Therapeutics, Waltham, MA) for supplying NO generators, accessory equipment and trouble-shooting advice; Kelin Li and Jeffrey Aubé (University of North Carolina) for synthesizing GSK1867513A; Alfonso Mendoza-Losano, GSK, Tres Cantos, Spain for supplying the bromoindazole; Omar Vandal (Gates Biotechnology Accelerator, Seattle, WA), Khisi Mdluli (Gates Medical Research Institute, Boston, MA), David Hermann (Gates Foundation, Seattle, WA); Lorenzo Berra, Binglan Yu, and Ryan Carroll (Mass General Brigham, Boston, MA); Ben Gold (Weill Cornell Medicine) for advice; and Sebastián Carrasco and Ileana C. Miranda (Center of Comparative Medicine and Pathology, Weill Cornell Medicine and Memorial Sloan Kettering Cancer Center) for evaluation of histopathology. This work was supported by a grant from the Gates Biotechnology Accelerator and by the Milstein Program in Chemical Biology and Translational Medicine. The Department of Microbiology and Immunology is supported by the William Randolph Hearst Trust.

## AUTHOR CONTRIBUTIONS

XJ performed the experiments and prepared the figures. CN designed the experiments, assisted with methemoglobin measurements, reviewed the literature and wrote the paper. The authors have no conflicts of interest.

