## Supplemental figures for "Effects of Exogenous Nitric Oxide Gas on *Mycobacterium tuberculosis* in vitro and in mice"

**A**

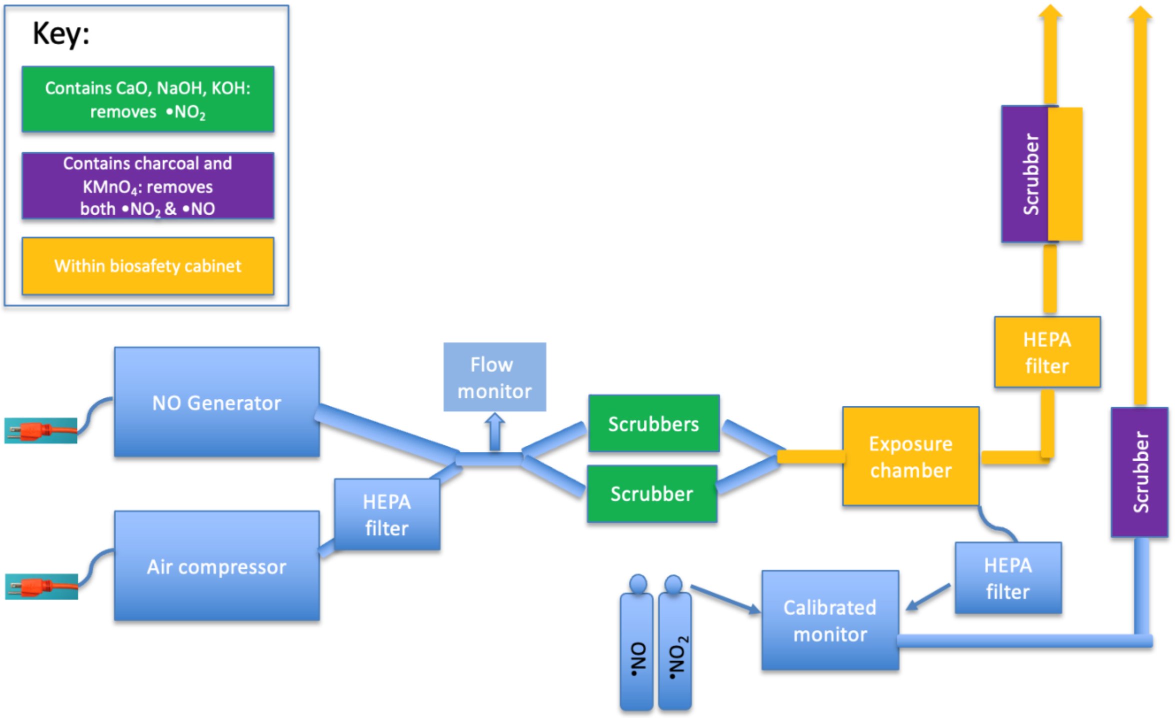
Key:
Contains CaO, NaOH, KOH: removes •NO2
Contains charcoal and KMnO4: removes both •NO2 & •NO
Within biosafety cabinet
NO Generator
Air compressor
HEPA filter
Flow monitor
Scrubbers
Scrubber
Exposure chamber
HEPA filter
Scrubber
Scrubber
HEPA filter
Calibrated monitor
•NO
•NO2

**B**

**C**

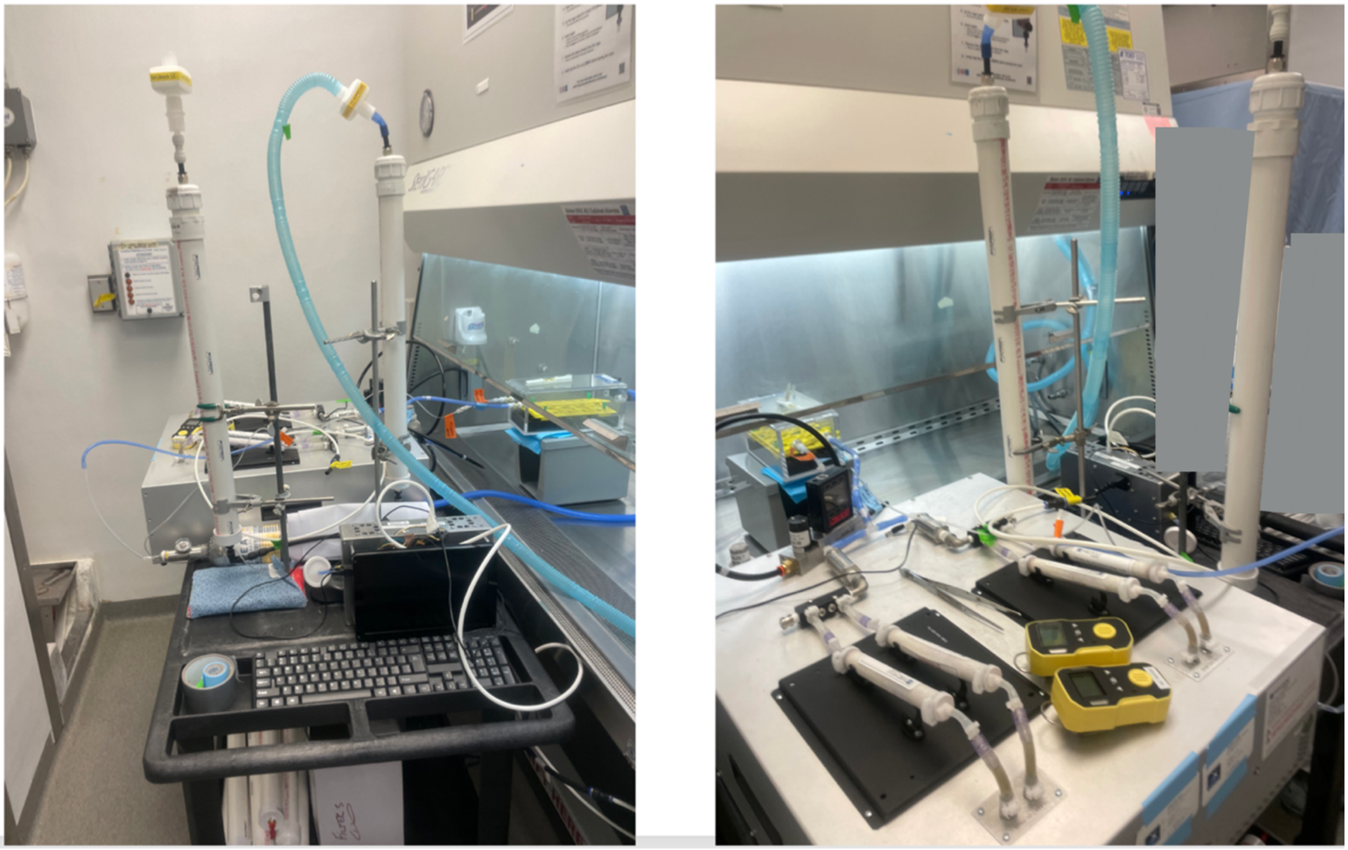

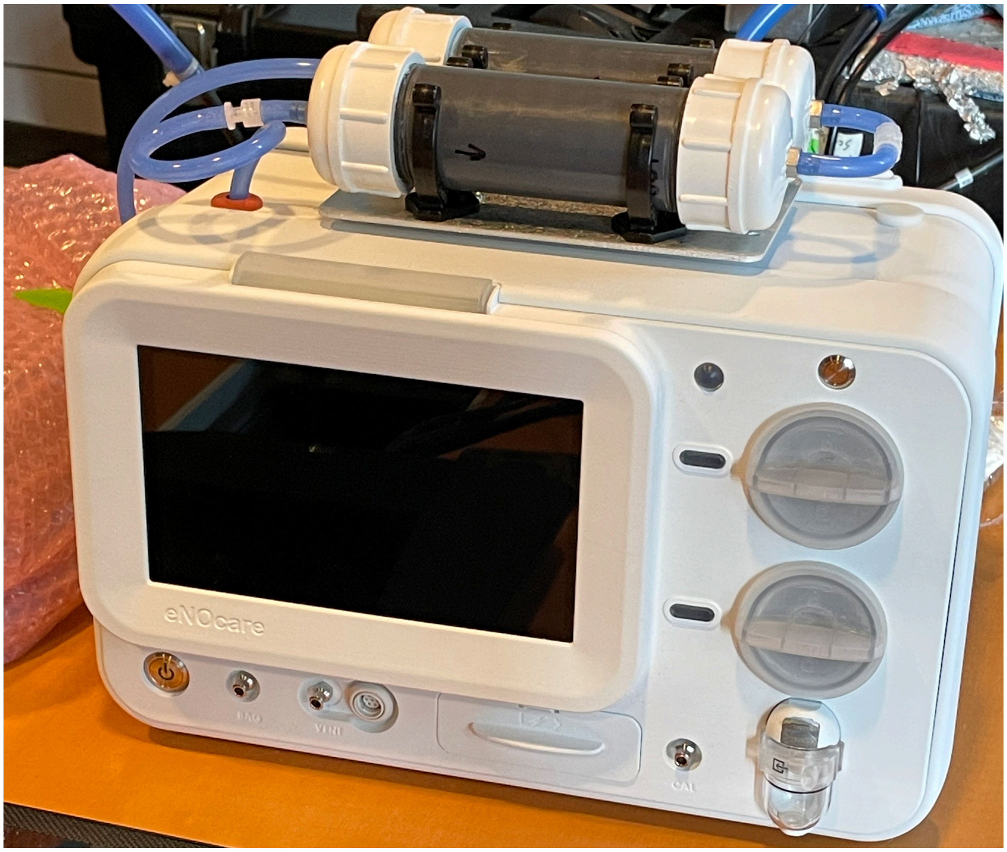


**Supplemental Figure 1. NO delivery system in a biosafety level 3 facility. (A) Schematic. (B) Custom-built unit. Clinical prototype.** Equipment shown in **(B)** and **(C)** was a kind gift of Third Pole Therapeutics.

A

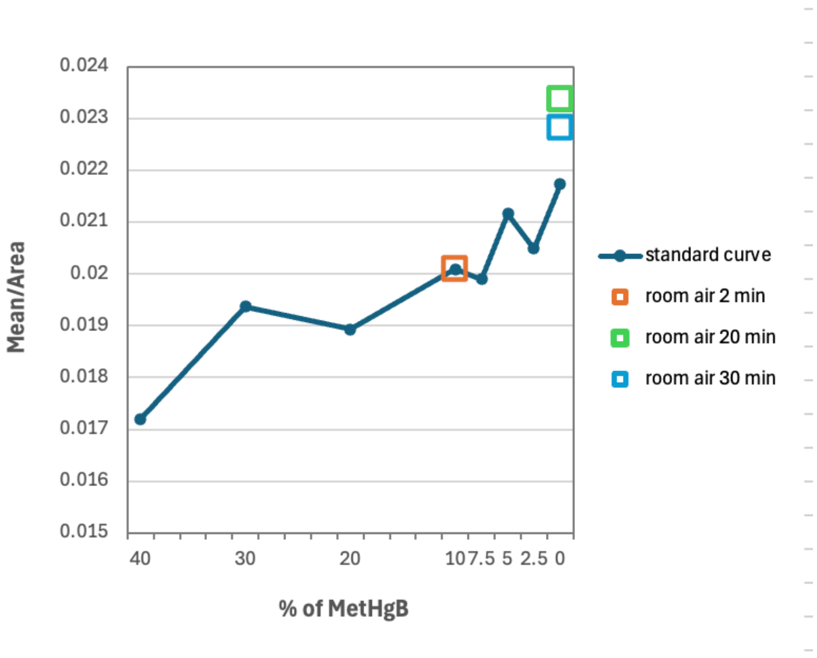

0.024
0.023
0.022
0.021
0.02
0.019
0.018
0.017
0.016
0.015
Mean/Area
40
30
20
10
7.5
5
2.5
0
% of MetHgB
standard curve
room air 2 min
room air 20 min
room air 30 min


B

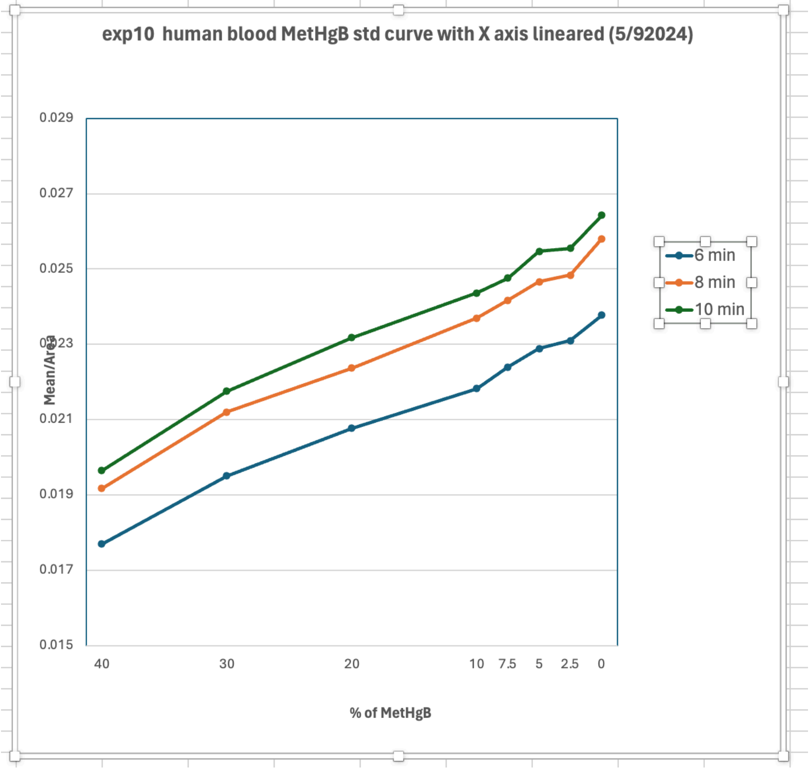

exp10 human blood MetHgB std curve with X axis lineared (5/92024)
0.029
0.027
0.025
0.023
0.021
0.019
0.017
0.015
Mean/Area
40
30
20
10
7.5
5
2.5
0
% of MetHgB
6 min
8 min
10 min

**Supplemental Figure 2. Estimation of methemoglobin level in blood of uninfected mice using a blood spot colorimetric method.** **(A)** Tail vein blood was collected and processed as quickly as possible (within 2 min) after 15 min exposure of mice to NO at 300 ppm and 20 and 30 min later. Methemoglobin levels were estimated by reference to a standard curve prepared in mouse blood (std). **(B)** Standard curves prepared in human blood with color red at indicated times after spotting on filters.
